# Inferring Accumulation Times of Mitochondrial DNA Deletion Mutants from Cross-Sectional Single-Cell Data: Application to Tabula Muris Senis

**DOI:** 10.64898/2026.01.30.702741

**Authors:** Axel Kowald, Thomas B L Kirkwood

## Abstract

Mitochondrial DNA (mtDNA) deletion mutants accumulate clonally in post-mitotic cells during ageing and can reach high intracellular fractions that impair oxidative phosphorylation. Despite extensive experimental documentation, the mechanisms underlying this accumulation and the timescales over which deletion mutants expand within individual cells remain poorly constrained. In a companion study, we developed a stochastic modelling framework based on a Moran birth-death process and showed that key parameters of mtDNA dynamics, mutation probability, selection advantage, and the fraction of advantageous mutations, can be inferred from cross-sectional single-cell RNA sequencing (scRNAseq) data. Here, we apply this framework to experimental scRNAseq data from the Tabula Muris Senis project, which provides a lifespan-resolved atlas of mouse tissues. We identify and quantify mtDNA deletions in limb muscle cells across multiple ages. Gene-resolved analyses reveal pronounced deletion hotspots associated with elevated heteroplasmy at ND5 and ND6, consistent with predictions that disruption of transcriptional feedback regulation confers a replication advantage. Conversely, deletions affecting ATP6 are associated with reduced heteroplasmy, suggesting an unexpected role of this gene in supporting mtDNA propagation. The distribution of unique deletion species per cell follows a Poisson distribution with a low mean, indicating limited intracellular mutant diversity, further constraining plausible accumulation mechanisms. Parameter fits yield selection advantage estimates that imply mean accumulation times of approximately 3 months from first occurrence to near-complete dominance. Our results support a model in which mtDNA deletion accumulation is driven by rare mutation events followed by rapid clonal expansion, and they demonstrate the potential of single-cell omics data to quantify fundamental parameters of mitochondrial ageing dynamics.

## Introduction

Mitochondria occupy a singular position in the biology of ageing: they generate most cellular ATP through oxidative phosphorylation, yet they also carry their own genome (mtDNA), which is exposed to intrinsic biochemical stressors and can acquire somatic mutations over the lifespan. Among the various classes of mtDNA lesions observed in mammalian tissues, large-scale deletion mutants have received particular attention because they can clonally expand within individual post-mitotic cells, especially neurons and skeletal muscle fibres, until they exceed functional thresholds for respiratory chain impairment^1-11^. This phenomenon is now supported by extensive single-cell and microdissection evidence across species and tissues, linking high mutant loads to cytochrome c oxidase (COX) deficiency, electron transport system (ETS) abnormalities, and progressive cellular pathology^12-15^.

A key conceptual challenge is that the tissue-level abundance of deletions in homogenates is often low, while individual affected cells can harbour near-homoplasmic mutant burdens. This spatial and cellular mosaicism is particularly clear in skeletal muscle, where ETS-abnormal regions appear segmentally within fibres and can progress to atrophy, splitting, breakage, and ultimately fibre loss, thereby contributing directly to sarcopenia^2,5,12-15^. In the nervous system, substantia nigra neurons show exceptionally high deletion burdens that frequently exceed the approximate threshold for respiratory deficiency, with strong correspondence between deletion load and COX deficiency in single neurons^1,3,6^. These observations establish mtDNA deletion accumulation as a conserved feature of mammalian ageing biology and motivate the central mechanistic question: by what process can a deleterious mutant genome reliably rise from rarity to dominance inside a cell?

Multiple explanatory frameworks have been proposed. Early “vicious cycle” ideas suggested that mitochondrial dysfunction elevates reactive oxygen species (ROS) production, accelerating further mtDNA mutagenesis and driving runaway deterioration^16,17^. However, a major empirical constraint is that many COX-deficient regions are dominated by a single deletion species rather than a broad mixture of independent mutants, which is difficult to reconcile with a strongly ROS-driven, continuously mutagenic process operating late in life^18^. Furthermore, as reviewed by^10^, there seems to be a surprising lack of mutations caused by ROS. Alternative proposals shifted the focus from mutation input to intracellular selection: “survival of the smallest” posits a replicative advantage for shorter genomes^19,20^, while “survival of the slowest” proposes that dysfunctional mitochondria evade turnover because they generate fewer damaging by-products and are degraded less efficiently^18,21^. Each of these ideas captures aspects of mitochondrial population dynamics, yet each also faces conceptual and experimental limitations. For instance, it is improbable that variations in genome size significantly affect replication speed^22^, and mechanisms of selective mitophagy seem to target dysfunctional mitochondria for removal rather than preservation.

A more parsimonious route to clonal dominance is neutral drift under relaxed replication. In this framework, mtDNA molecules turn over continuously; replication and degradation introduce stochastic fluctuations in heteroplasmy such that, given sufficient time and suitable effective population sizes, some mutants can expand to high levels without invoking positive selection^23,24^. This class of models can reproduce several features of human ageing data, including the existence of rare but high-burden mutant cells. Yet a critical cross-species constraint emerges when these models are confronted with short-lived animals. Parameter regimes that generate enough COX-deficient cells within a mouse or rat lifespan tend to predict a higher diversity of coexisting mutant species within individual cells than is typically observed in single-fibre and single-neuron studies^25^. This tension suggests that, at least in some contexts, additional selective structure may be required to explain why affected cells often harbour one dominant deletion species rather than many.

In the first manuscript of this two-part work^26^, we addressed this mechanistic impasse by combining biological reasoning with mathematical modelling and by developing an inference framework that extracts two quantities that are otherwise experimentally inaccessible at single-cell resolution, namely the selection advantage of deletion mutants and the distribution of their intracellular accumulation times. The key methodological insight is that although direct longitudinal tracking of mtDNA genotypes in single cells is impossible (because measuring genotype destroys the cell), cross-sectional single-cell RNA sequencing (scRNAseq) provides a practical surrogate since mtDNA deletions that remove gene segments also produce mitochondrial transcripts with corresponding deletions. Consequently, the transcriptome can encode quantitative information about the underlying mtDNA composition, allowing mutant-to-wild-type ratios to be estimated indirectly from observed expression patterns. Using synthetic scRNAseq-like data, generated via stochastic models of the mitochondrial life-cycle, and analysing these data with a Moran birth–death process, we showed that selection advantage, mutation probability, and the fraction of mutants that display a selection advantage can be recovered from cross-sectional snapshots. Once the selection advantage is known, the same Moran framework yields distributions of mutant accumulation times from first appearance to functional takeover.

The present manuscript represents the second part of the overall work and moves from proof-of-principle to experimental application. Here, we apply the established mathematical and bioinformatic pipeline to scRNAseq data from the Tabula Muris Senis (TMS) project^27^, which provides a lifespan-resolved atlas of mouse tissues across multiple ages. TMS offers two advantages that are directly aligned with the central questions of mtDNA deletion dynamics. First, it provides large numbers of single-cell transcriptomes, enabling detection and quantification of rare subpopulations that may correspond to cells at different stages of mutant takeover. Second, it samples multiple age groups, allowing cross-sectional “snapshots” to be arranged along a temporal axis in a way that is compatible with our modelling approach. Together, these properties make TMS an unusually powerful dataset for estimating parameters governing in vivo mtDNA mutant accumulation in a mammalian model.

Our primary objective in this manuscript is to obtain quantitative estimates of intracellular accumulation times for mitochondrial deletion mutants from real experimental data. This objective is not merely descriptive. Accumulation time is a discriminating parameter since different mechanistic hypotheses imply qualitatively different temporal regimes. In particular, models in which drift dominates without strong selection tend to predict broad, age-dependent distributions strongly shaped by early-life founder events, whereas models with a genuine selective advantage for particular deletion classes predict a biphasic dynamic in which long waiting times for the right mutation are followed by comparatively rapid takeover once such a mutant arises. The transcription-coupled replication hypothesis of Kowald and Kirkwood^28,29^ provides a concrete molecular route to such a selective advantage. Because mitochondrial replication initiation is linked to transcription, deletions that remove components of a negative feedback loop can become persistently over-transcribed and thereby gain a cis-acting replication advantage. Strikingly, surveys of clonally expanded deletions across species indicate strong enrichment for deletions overlapping ND4 (often also ND5), consistent with disruption of a conserved feedback architecture^28,29^. Under this view, the relevant biological question becomes not only whether deletions are present, but whether their inferred selection advantage and takeover times are compatible with the observed incidence of mitochondrial dysfunction across ages in mice.

By anchoring the analysis in experimentally observed single-cell heterogeneity, and by translating that heterogeneity into mechanistically interpretable parameters, we aim to provide quantitative constraints on competing explanations for mitochondrial mutant accumulation. These constraints, in turn, should help distinguish whether the observed age-associated rise in respiratory-deficient cells is consistent with predominantly neutral drift, requires positive selection for a subset of deletion classes, or reflects a combination of stochastic mutation input and selective amplification shaped by mitochondrial regulatory architecture.

## Results

### The Tabula Muris Senis Dataset

The Tabula Muris Senis dataset constitutes a comprehensive and systematically designed single-cell transcriptomic resource aimed at characterizing the molecular and cellular manifestations of ageing across the murine lifespan^27^. By resolving transcriptomes at single-cell resolution across a broad age range, Tabula Muris Senis provides a uniquely suitable framework for analysing stochastic and heterogeneous processes, such as the accumulation of mitochondrial DNA (mtDNA) deletion mutants, that operate at the level of individual cells.

In total, single-cell RNA sequencing (scRNA-seq) was performed on more than 350000 cells isolated from 23 distinct tissues and organs of male and female C57BL/6JN mice. Animals were sampled at six ages (1, 3, 18, 21, 24, and 30 months) spanning early postnatal development, young adulthood, middle age, and extreme old age (Fig. 1A & B). Two complementary scRNA-seq technologies were employed: a high-throughput droplet-based platform (10x Genomics) to capture large numbers of cells, and a plate-based Smart-seq2 approach.

**Fig. 1.**
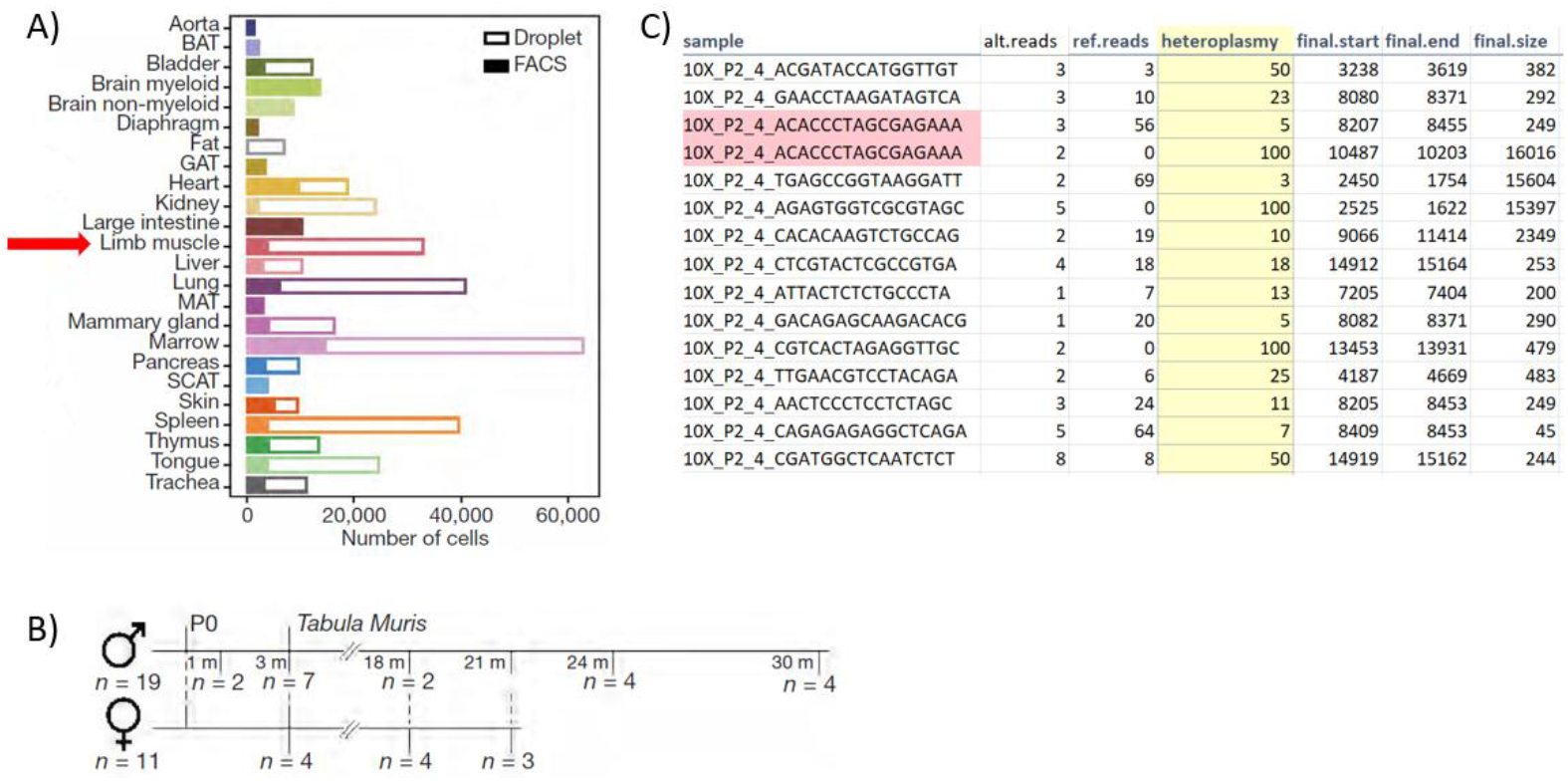
Overview of the Tabula Muris Senis dataset and extracted data. A) 23 different tissues from male and female mice were analysed using FACS and microfluidic droplets. We used post-mitotic limb muscle cells for our analysis. B) Measurements were performed at six different ages ranging from 1 month to 30 months. However, limb muscle data were only available starting from 3 months of age. C) Example output from our software MitoSAltsc, which finds and extracts cells containing mRNA reads with deletions (here for limb muscle cells at the age of 30 months). The entry highlighted in red stems from a single cell harbouring two different deletion mutants with heteroplasmy values of 5% and 100%. See main text for further details.

The availability of data from multiple age groups is a critical prerequisite for the analysis presented in this work. Our parameter estimation strategy relies on fitting a stochastic Moran birth–death process to age-resolved curves derived from cross-sectional data^26^. Such an approach requires several temporally spaced sampling points to constrain the underlying dynamics and to distinguish between differences in mutation occurrence, selection strength, and accumulation time distributions. The Tabula Muris Senis dataset satisfies this requirement in an unusually comprehensive manner, providing sufficiently dense age coverage to enable meaningful inference of time-dependent processes from static observations.

Furthermore, the dataset includes a broad range of terminally differentiated and post-mitotic cell types, which is essential given that mtDNA deletion mutants are known to accumulate preferentially in cells with limited or absent proliferative capacity. In proliferating cells, clonal expansion of mtDNA variants is continuously diluted by cell division, whereas in post-mitotic cells mutant genomes can persist and expand intracellularly over long timescales. For this reason, we restrict our analysis to limb skeletal muscle, a tissue in which post-mitotic myofibers constitute the dominant cell population and in which age-associated clonal expansion of mtDNA deletions has been extensively documented. Within the Tabula Muris Senis dataset, limb muscle transcriptomic data are available for animals aged 3, 18, 21, 24, and 30 months, providing five age points that span the majority of adult and late-life stages relevant for mitochondrial mutant accumulation.

To identify and quantify mitochondrial DNA deletion events at single-cell resolution, we created MitoSAltsc, a version of the MitoSAlt pipeline (originally developed for accurate identification of mtDNA deletions and duplications from deep genomic sequencing data^30^) that is compatible with scRNAseq data (see Methods). This software enables detection of split-read signatures indicative of mtDNA deletions and provides cell-resolved estimates of deletion breakpoints and heteroplasmy. In the majority of cases, a single deletion species is detected per cell, consistent with previous observations that clonally expanded mtDNA deletions often dominate within individual post-mitotic cells. However, a subset of cells contains more than one deletion event, each with its own heteroplasmy level. Such cases, highlighted in red in Fig. 1C, demonstrates the ability of the pipeline to resolve multiple coexisting structural variants within a single cell and provide important information about the diversity of mutant species.

The combination of the Tabula Muris Senis dataset and the MitoSAltsc pipeline provides a powerful experimental and computational foundation for quantitative analysis of mtDNA deletion dynamics. The age-resolved, single-cell nature of the data enables inference of accumulation processes that unfold over months to years, while the ability to accurately detect and quantify deletion heteroplasmy at single-cell resolution is essential for linking observed transcriptomic patterns to underlying mitochondrial population dynamics.

### Deletion Hotspots

In earlier theoretical and computational work, it was proposed that only a specific subset of mitochondrial DNA deletions should experience a genuine intracellular selection advantage, namely those that disrupt a transcriptional feedback mechanism linking mitochondrial gene expression to replication initiation^28,29^. Consistent with this hypothesis, analyses of single-cell and single-fibre datasets from mice, rats, rhesus monkeys, and humans have repeatedly shown that the vast majority of clonally expanded deletions overlap genes encoding subunits of respiratory chain complex I, most notably ND4 and, frequently, ND5.

The Tabula Muris Senis dataset offers a new and complementary opportunity to test this hypothesis in an unbiased, genome-wide manner and at single-gene resolution. Unlike earlier studies that relied on manually curated collections of clonally expanded deletions, the present analysis leverages large-scale single-cell transcriptomic data to systematically assess whether deletions affecting specific mitochondrial genes are associated with higher or lower levels of heteroplasmy. To this end, we stratified cells based on whether a given mitochondrial gene was affected by a deletion or not, and compared the resulting heteroplasmy distributions between the two groups. For each protein-coding mitochondrial gene, and separately for each age class, we performed a two-tailed t-test contrasting the heteroplasmy levels of the two groups. The resulting p-values were transformed by taking the negative logarithm to facilitate comparison across genes and ages, and these values are displayed in Fig. 2.

**Fig. 2.**
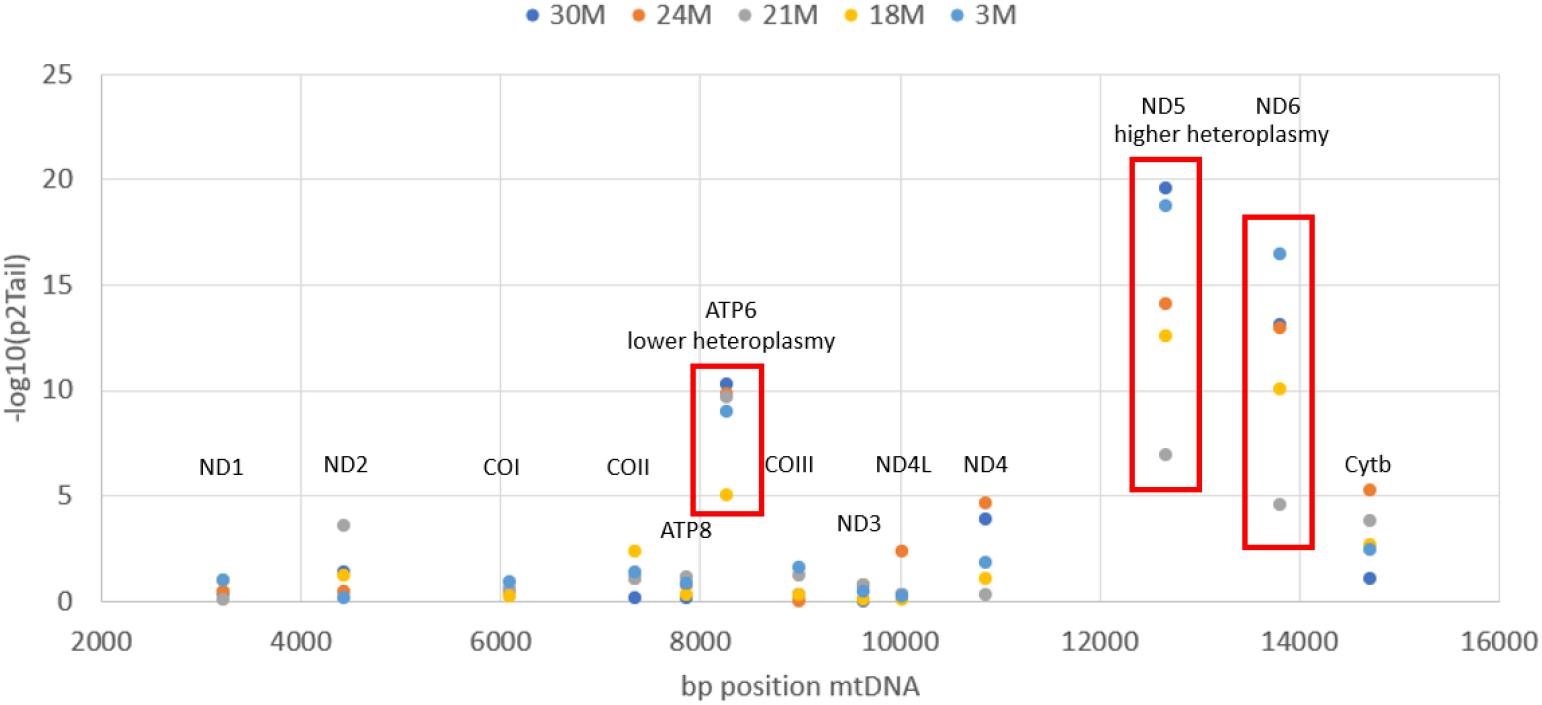
Two-tailed t-test p-values for all mitochondrial protein-coding genes across the different age classes, comparing heteroplasmy levels between cells in which a given gene is affected by a deletion and cells in which it is not. For improved visualization and comparability, p-values are displayed as their negative logarithm. Red boxes indicate the three genes that exhibit exceptionally low p-values, reflecting particularly strong and consistent differences in heteroplasmy associated with deletions affecting these loci.

Most mitochondrial genes show only modest or negligible differences in heteroplasmy between affected and unaffected cells, with negative log p-values close to or below unity across ages. In contrast, three genes stand out with exceptionally strong statistical signals: ATP6, ND5, and ND6. For these genes, negative log p-values reach values between approximately 10 and 20, far exceeding those observed for any other gene and indicating highly significant differences in heteroplasmy. Importantly, these effects are not confined to a single age class but are reproducible across multiple ages, suggesting that they reflect intrinsic properties of the affected genes rather than age-specific sampling artefacts.

To interpret the directionality of these effects, we performed additional one-tailed t-tests. This analysis revealed a clear dichotomy between the genes involved. For ND5 and ND6, deletions affecting these genes are associated with significantly higher levels of heteroplasmy compared with deletions that spare them. This result is fully consistent with the transcriptional feedback escape hypothesis and would indicate that loss of ND5 or ND6 confers a replication advantage that allows mutant genomes to expand to higher intracellular fractions. The effect is particularly pronounced for ND5, which exhibits the strongest and most consistent signal across all age groups, in line with previous observations.

In contrast, the situation for ATP6 is markedly different. Although deletions affecting ATP6 also yield extremely low p-values, the direction of the effect is opposite, implying that cells in which ATP6 is disrupted by a deletion tend to exhibit lower, rather than higher, levels of heteroplasmy. This finding indicates that ATP6 deletions are selectively disadvantaged, either because they impair replication, transcription, or mitochondrial maintenance in a way that limits clonal expansion. This unexpected result highlights that not all deletions affecting coding regions are equivalent and that some genes may play an active, positive role in supporting the transcription and replication machinery. We return to this point in the discussion section, where alternative mechanistic explanations are considered.

Taken together, these results provide strong, independent support for the central idea that only a subset of mitochondrial deletions, specifically those affecting ND5 and ND6, enjoy a selective advantage that promotes clonal expansion. At the same time, the data refine the original hypothesis by indicating that other genes, such as ATP6, may be essential for successful mutant proliferation and thus act as negative determinants of heteroplasmy when deleted. Beyond their mechanistic implications, these findings have an important quantitative consequence for our modelling framework. They allow us to estimate *fracAdv*, the fraction of all deletion mutations that confer a selective advantage. Given the total length of the murine mitochondrial genome (approximately 16.3 kb) and the combined genomic span of the ND5 and ND6 coding regions, we estimate *fracAdv* to be on the order of 0.14. This estimate substantially constrains the parameter space of our stochastic Moran model. Previous analyses using synthetic data have shown that *fracAdv* is more difficult to infer robustly than the mutation probability (*mutProb*) or the selection advantage (*selAdv*) ^26^. By fixing *fracAdv* to an empirically grounded value, we reduce the dimensionality of the fitting problem from three parameters to two, thereby improving identifiability and stability of the subsequent inference of accumulation times from experimental data.

### Distribution of unique mutants

The number of distinct mitochondrial deletion species present within a single cell provides important information about the underlying mechanisms governing mutant accumulation. In particular, it allows one to discriminate between scenarios dominated by stochastic mutation input and drift, which tend to generate multiple coexisting mutant lineages, and scenarios in which strong intracellular selection favours the expansion of a small subset of advantageous mutants, leading to low intracellular diversity. For this reason, the distribution of unique mutant types per cell constitutes a sensitive and conceptually informative summary statistic of mitochondrial population dynamics and plays a central role in the modelling framework developed in the first part of this work.

In the simulation study presented in part one^26^, we showed that under a broad range of biologically plausible assumptions, the number of unique mutant species per cell is well described by a Poisson distribution. This result follows naturally from the fact that deletion events arise stochastically and independently over time, while only a subset of these events survives long enough to be detected in a cross-sectional snapshot. In that context, the single parameter of the Poisson distribution, λ, represents the mean number of distinct mutant species per cell and provides a compact measure of intracellular mutant diversity. Importantly, our simulations further demonstrated that λ is not an arbitrary quantity but is systematically shaped by age as well as the underlying model parameters. Depending on the balance between mutation probability, selection advantage, and the fraction of advantageous mutations, λ was predicted to increase approximately linearly or quadratically with age, reflecting the cumulative opportunity for successful mutant lineages to arise and persist.

Fig. 3A-E shows that the experimental data from the Tabula Muris Senis limb muscle cells closely mirror this key qualitative prediction in that the empirical distributions of the number of unique mutant species per cell at each age are well approximated by Poisson distributions. This agreement provides an important validation of the modelling assumptions and indicates that, despite the complexity of the biological system and the limitations of transcriptomic data, the dominant statistical structure of mutant diversity is captured by the same stochastic principles identified *in silico*. The fitted values of the Poisson parameter λ are all of order 0.1, consistent with the notion that most cells harbour either no detectable deletion or a single dominant mutant species, with only a small fraction containing more than one distinct deletion. This low level of intracellular diversity is incompatible with purely neutral drift operating over the entire lifespan and instead supports the presence of selective or regulatory constraints that limit the coexistence of multiple mutant lineages within a single cell.

**Fig. 3.**
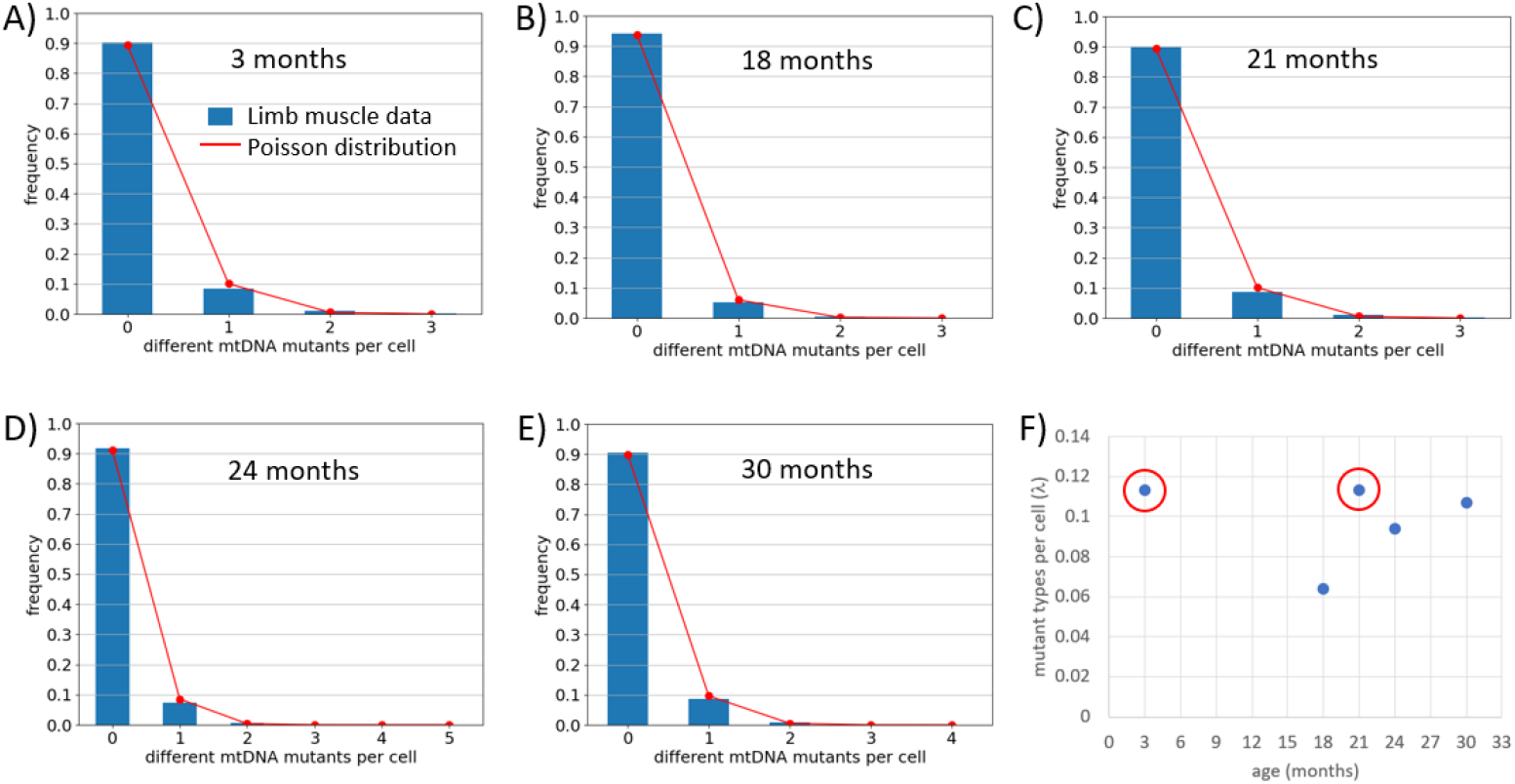
A) to E) Distributions of the number of unique deletion mutants per limb muscle cell at different timepoints during a mouse lifespan. A value of zero indicates cells containing only wild-type mtDNAs. Blue bars show the experimental results and red lines indicate the corresponding Poisson distributions. The number of analysed cells range from 3285 to 7178. F) Dependence of the mean number of unique mutants depending on age. Red circles indicate data points that may be outliers.

When λ is plotted as a function of age (Fig. 3F), however, the experimental data reveal a more nuanced picture than suggested by the simulations. While the overall magnitude of λ closely matches that obtained in the synthetic data, the age dependence does not follow a simple monotonic linear or quadratic trend. Instead, the values fluctuate across ages, with comparatively higher estimates at 3 and 21 months. One possible interpretation is that the points at 3 and 21 months represent statistical outliers, in which case the remaining values would be broadly consistent with the expected age dependence derived from the model. Such deviations are not unexpected given the characteristics of the dataset. The number of limb muscle cells available at each age is limited, and the effective detection of deletion species relies on a relatively small number of mitochondrial transcripts per cell. Both factors can introduce substantial sampling noise into estimates of rare events such as low-heteroplasmy deletions and can therefore distort the inferred value of λ. We defer a more detailed interpretation of the non-monotonic age dependence of λ to the Limitations and Discussion section, where these observations and their implications for parameter inference are addressed in greater depth.

### Obtaining Estimates for *selAdv* and *mutProb*

In the previous section, we noted that the data points corresponding to 3 and 21 months of age deviate from the pattern observed at the other ages. A similar effect becomes apparent when we analyse the fraction of cells with more than a given fraction of mitochondrial mutants, which constitutes the central observable used for parameter inference in our modelling framework (Fig. 4A; see also^26^). These curves, denoted fcwm00, fcwm05, fcwm20, and fcwm50, describe the age-dependent fraction of cells exceeding 0%, 5%, 20%, and 50% mutant load, respectively, and serve as the empirical targets for fitting the parameters of the stochastic Moran birth–death process introduced in part one.

**Fig. 4.**
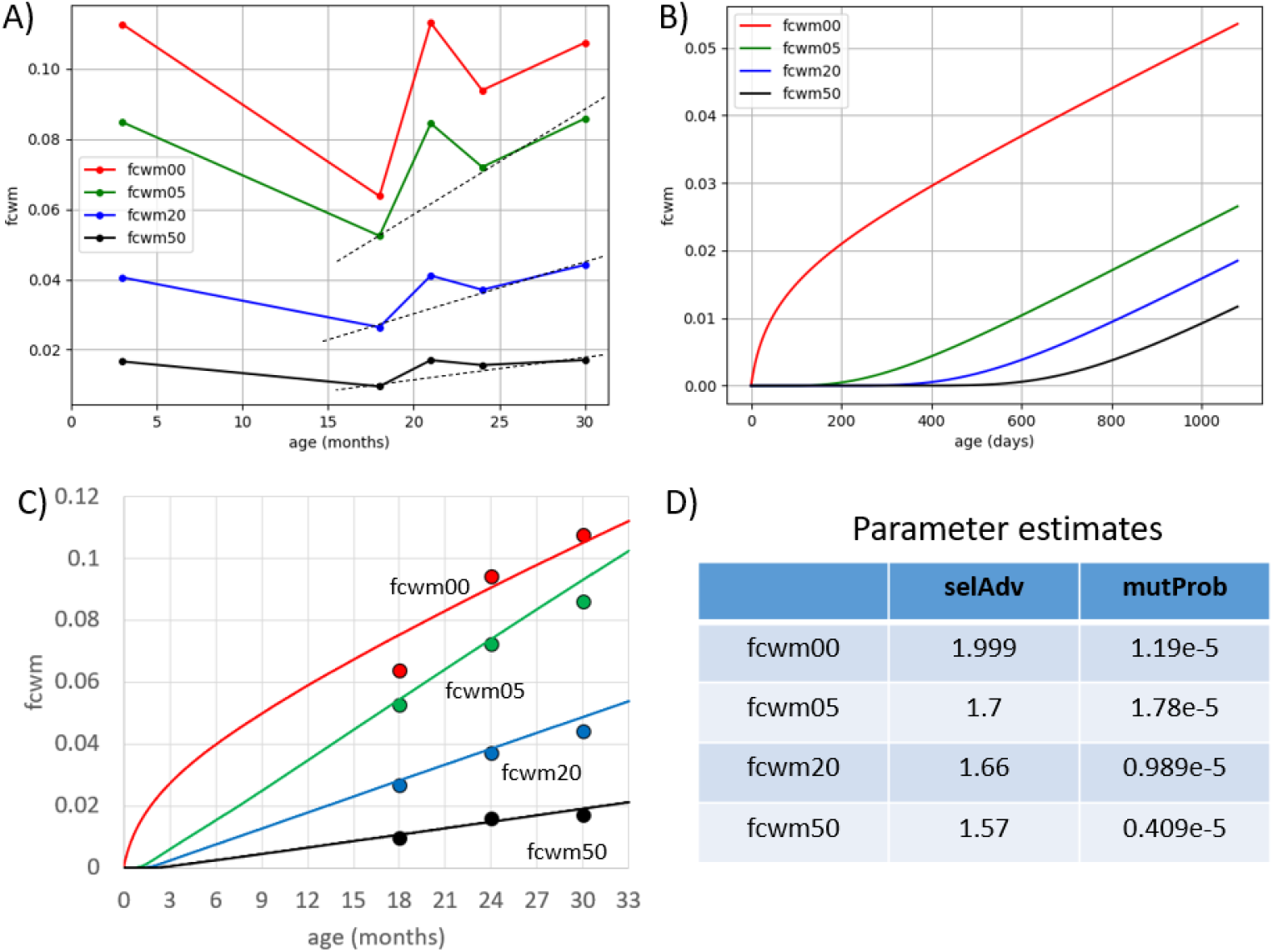
A) Fraction of cells exceeding 0%, 5%, 20%, and 50% mutant mtDNA, derived from the Tabula Muris Senis dataset. Dotted lines indicate slopes for fcwm05, fcwm20, and fcwm50, excluding the 3- and 21-month data points. B) Reproduction of Fig. 6B from^26^, illustrating the characteristic behaviour of fcwm curves generated by the Moran model used in the simulation study. C) Independent fits of the Moran model to individual fcwm levels for parameter estimation. D) Resulting mean estimates of *mutProb* and *selAdv* for the different fcwm thresholds based on 40 repetitions of the fitting procedure (see main text for details).

Briefly, the Moran-based approach models the intracellular mtDNA population as a fixed-size pool subject to stochastic replication and degradation events^31-33^. Mutant genomes arise with probability *mutProb* and may enjoy a selective replication advantage *selAdv* relative to wild type. For a given parameter set, simulations yield age-dependent curves for the fraction of cells exceeding specified mutant thresholds, which can be directly compared to the experimental fcwm curves. In the simulation study presented in part one, all fcwm curves exhibited similar slopes and differed primarily by horizontal shifts, reflecting different threshold levels being crossed at different times.

Fig. 4B shows a representative example of such simulated fcwm curves. Comparison with the experimental curves in Fig. 4A reveals two systematic differences. First, in the simulated data there is a pronounced separation between the curves for fcwm00 and fcwm05, whereas in the experimental data these two curves lie much closer together. Second, the simulated curves exhibit nearly identical slopes, while in the experimental data the slopes differ across thresholds. Excluding the 3- and 21-month data points, the experimental curves display a clear ordering in which the slope decreases with increasing mutant threshold, such that fcwm05 increases more steeply with age than fcwm20, which in turn is steeper than fcwm50.

These discrepancies indicate that the assumptions underlying the simple Moran process, most notably homogeneous turnover and the absence of explicit quality control or state-dependent effects, do not fully capture the behaviour observed *in vivo*, or alternatively that limitations in data quantity and detection sensitivity distort the inferred curves. Possible biological explanations, including selective removal of low-level or deleterious mutants and heterogeneity in mutant fitness, are discussed in more detail in later sections. At this stage, the key practical implication is methodological.

Because the fitting strategy employed in the simulation study relies on the parallelism of fcwm curves across thresholds, it cannot be directly applied to the experimental data. Instead of performing a single joint fit to all fcwm curves simultaneously, we therefore fit the model separately to each fcwm level. This procedure yields four independent parameter estimates rather than a single consensus estimate. While this approach has clear limitations, it allows us to extract threshold-specific information from the data and to assess the consistency of inferred parameters across mutant load levels. The consequences and validity of this strategy are addressed further in the Discussion.

As shown in Fig. 4D fitting was performed for the parameters *selAdv* and *mutProb*, with the fraction of advantageous mutations, *fracAdv*, fixed at 0.14 based on the analysis shown above (section Deletion Hotspots). To minimize stochastic effects of the parameter estimation process, we repeated the process 40 times and report the mean of the obtained values. Across increasing fcwm thresholds, the inferred selection advantage is very similar, while the estimated mutation probability decreases. The fit based on fcwm00 constitutes a notable exception: it drives *selAdv* to the upper bound of the allowed fitting range (restricted to values between 0 and 2). Given the unusually small separation between fcwm00 and fcwm05 in the experimental data, and the strong sensitivity of fcwm00 to low-level and potentially transient mutant lineages, this result suggests that the fcwm00 curve violates key assumptions of the Moran-based model. Consequently, in the subsequent analysis we restrict attention to parameter estimates derived from fcwm05, fcwm20, and fcwm50.

### Accumulation times of deletion mutants

Using the Tabula Muris Senis data in combination with our analysis pipeline, we obtained empirical estimates for three key parameters governing mitochondrial mutant dynamics: the fraction of mutations that confer a selective advantage (*fracAdv*), the strength of that advantage (*selAdv*), and the mutation probability (*mutProb*). While all three parameters are required for a complete description of the stochastic process underlying mtDNA turnover, the selection advantage plays a particularly central role because it directly determines the timescale over which a deletion mutant, once it arises, can expand and ultimately dominate the intracellular mtDNA population. For this reason, this section focuses on translating the inferred values of *selAdv* into quantitative estimates of accumulation times.

The connection between selection advantage and accumulation time is established through the Moran birth and death process that forms the basis of our modelling framework. In this model, a cell contains a fixed number of mtDNA molecules that undergo continuous turnover through replication and degradation events. When a mutant genome enjoys a selective advantage, *selAdv*, its probability of being chosen for replication is increased relative to wild type. Starting from the appearance of a single mutant molecule, the Moran process defines a stochastic trajectory for the mutant fraction, which may either go extinct or expand toward fixation. Conditional on successful expansion, the model yields a distribution of first-passage times, defined as the time required for the mutant fraction to cross a specified threshold.

As described in detail in the section “Using the Moran process to model accumulation times” in part one^26^, these first-passage times can be calculated by simulating the Moran process with parameters fixed to the experimentally inferred values. For each simulation run, a single advantageous mutant is introduced into an otherwise wild-type population, and the process is followed until the mutant fraction either reaches a predefined dominance threshold or is lost. Repeating this procedure many times yields a distribution of accumulation times conditioned on successful takeover. In the present analysis, we define takeover operationally as reaching 90% mutant load, which provides a practical proxy for near-complete dominance while avoiding the long tails associated with exact fixation.

Fig. 5 shows the resulting distributions of accumulation times obtained using the estimate of *selAdv* derived in the previous section for fcwm20. As expected from the underlying dynamics of the Moran process, higher selection advantage leads to faster accumulation. For the largest estimated advantage (*selAdv* = 1.7), the mean accumulation time is approximately 96.2 days, whereas for the smallest advantage (*selAdv* = 1.57) the mean increases to about 101.2 days. The intermediate estimate (*selAdv* = 1.66) yields a mean accumulation time that lies between these two values.

**Fig. 5.**
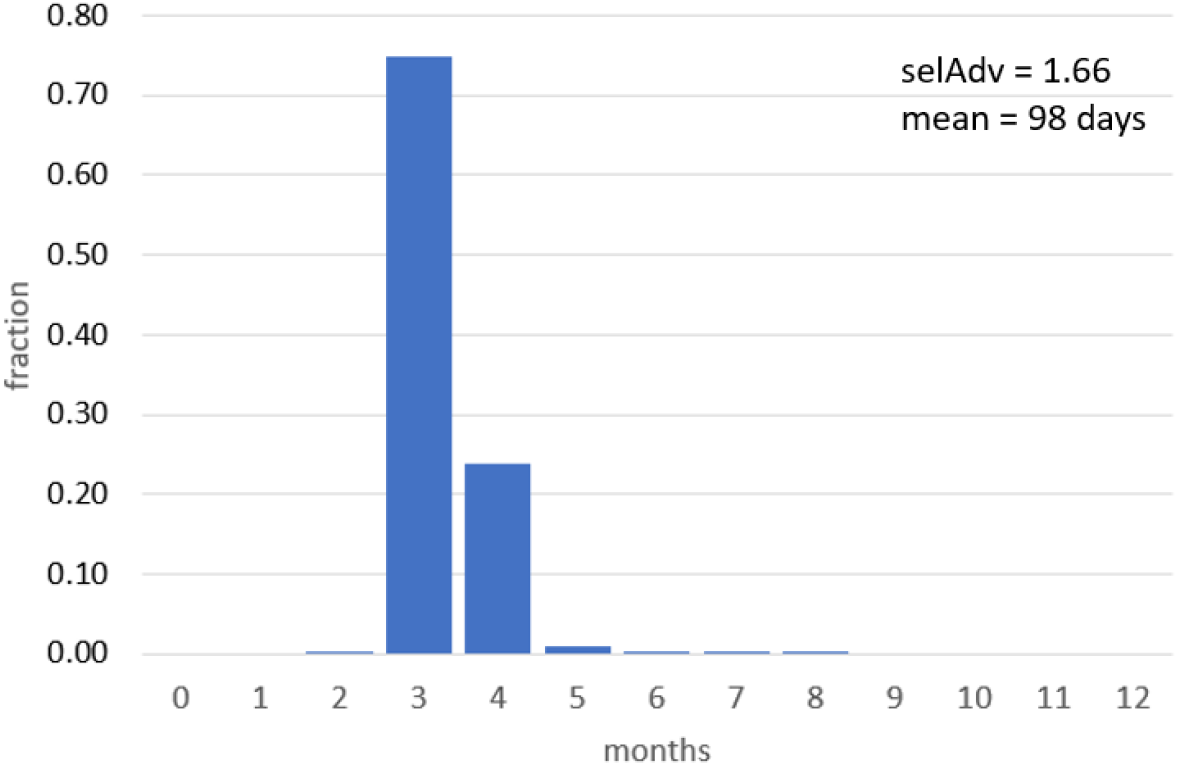
Distribution of accumulation times for mitochondrial deletion mutants (from first occurrence until 90% prevalence) based on an estimated parameter value for selection advantage *selAdv* of 1.66 (fcwm20). The selection advantages for fcwm05 and fcwm50 are very similar (see Fig. 4D) and also result in very similar mean accumulation times of 96.2 and 101.8 days.

Although the fitting procedure did not yield a single, unambiguous estimate for *selAdv*, the resulting accumulation times are remarkably consistent. Across all three parameter sets, the mean time required for a deletion mutant to expand from first occurrence to near-complete dominance is very fast and differs by only a few days. This narrow range indicates that, once an advantageous deletion arises, clonal expansion is predicted to be rapid on the timescale of the murine lifespan. Consequently, variability in the age at which mutant-dominated cells appear is likely driven primarily by the waiting time for the initial occurrence of a suitable deletion, rather than by large differences in expansion kinetics once such a mutant is present.

## Limitations

To estimate mutation probability and selection advantage, we employed a stochastic Moran model describing birth, death, and mutation of mtDNA molecules within individual cells^31-33^. While this framework captures key features of intracellular mtDNA dynamics and proved effective in the simulation study presented in part one, its agreement with the experimental results obtained from the Tabula Muris Senis (TMS) scRNAseq data is only partial (Fig. 4). In particular, the model predicts a substantially larger separation between the fraction of cells containing any mutant mtDNA (fcwm00) and those exceeding 5% mutant load (fcwm05) than is observed experimentally. In addition, whereas the model yields fcwm curves with nearly identical slopes across mutant thresholds, the experimental data indicate threshold-dependent slopes, with progressively shallower increases at higher mutant fractions. Finally, the data points corresponding to 3 and 21 months of age appear inconsistent with the overall trend defined by the remaining age classes (Fig. 3, Fig. 4).

One likely contributor to these discrepancies is the limited amount of mitochondrial information contained in standard scRNAseq datasets. In the TMS data, the number of reads mapping to mitochondrial genes is very small: e.g. for the 30 months age class, the maximum number of wild-type mitochondrial reads per cell is 433, with a mean of only 20.5, while reads containing deletions did not exceed 10 per cell and average around 2.1. As a consequence, estimates of heteroplasmy at the single-cell level are inherently noisy and subject to substantial sampling error. This uncertainty propagates directly into the calculation of fcwm values, particularly at low mutant fractions where classification depends on very few reads. Experimental strategies that enrich for mitochondrial transcripts, or sequencing protocols optimized for higher mitochondrial coverage, would substantially reduce this source of uncertainty and allow more reliable inference of mutant fractions.

A second, conceptually distinct limitation is the simplicity of the underlying Moran model. The model assumes homogeneous turnover dynamics and does not include explicit mitochondrial quality control mechanisms. In living cells, processes such as mitophagy are known to preferentially remove dysfunctional mitochondria and may act more strongly at low to intermediate heteroplasmy levels. Such selective clearance would reduce the lifetime of deleterious or neutral mutants shortly after their appearance, thereby depleting the pool of transient low-level mutants. This effect would naturally bring fcwm00 closer to fcwm05, as cells with any detectable mutants would be enriched for lineages capable of partial expansion. Moreover, the model assumes that replication dynamics are independent of the current level of heteroplasmy. If replication, degradation, or quality control processes depend on mutant load, for example through declining ATP production, altered membrane potential, or activation of stress responses, then expansion kinetics would slow as heteroplasmy increases. This could explain the experimentally observed ordering of slopes, with fcwm05 rising more steeply than fcwm20 and fcwm50. While incorporating such effects would increase biological realism, it would also add complexity and additional parameters. Given the current data limitations, it is not yet clear whether such model extensions are warranted or whether improved data quality alone would reconcile model and experiment.

A further limitation arises from the use of mitochondrial transcripts as a proxy for mitochondrial genomes. If, as proposed previously^28,29^, certain deletions disrupt transcriptional feedback inhibition and thereby increase transcriptional output, then deletion-bearing mtDNA molecules may be overrepresented at the RNA level relative to their true genomic abundance. In this case, heteroplasmy estimates derived from mRNA would systematically overestimate mtDNA frequencies for advantageous mutants. Single-cell DNA sequencing (scDNAseq) would circumvent this problem by directly quantifying mtDNA copy number and deletion frequencies at the genomic level. In addition, scDNAseq would allow direct measurement of total mtDNA copy number per cell, which is assumed to be constant (1000 genomes) in the present work. This assumption is a strong simplification, as mtDNA copy number is known to vary across cell types, ages, and physiological states. Future scDNAseq studies would therefore represent a major advance for quantitative analyses of mitochondrial population dynamics. Recently, scDNAseq techniques have been developed^34-36^ and it would be very useful if they would be applied to ageing studies.

## Discussion

The accumulation of mtDNA deletion mutants in post-mitotic cells is a well-established hallmark of ageing in mammals. Over the past decades, a range of mechanistic explanations has been proposed, spanning purely stochastic models based on random drift to hypotheses invoking selective advantages conferred by specific deletions, for example through disruption of transcriptional feedback regulation. Discriminating between these competing ideas requires quantitative estimates of key kinetic parameters, in particular the time required for a mutant to expand from its initial occurrence to dominance within a cell.

In a companion study, we demonstrated that such parameters, specifically the fraction of advantageous mutations (*fracAdv*), the mutation probability (*mutProb*), and the selection advantage (*selAdv*), can be inferred from cross-sectional single-cell transcriptomic data using a stochastic Moran framework^26^. In the present work, we applied this analysis pipeline to experimental scRNAseq data from the Tabula Muris Senis project, thereby testing the feasibility of extracting biologically meaningful accumulation times from real ageing data.

The TMS dataset revealed clear gene-specific patterns in deletion-associated heteroplasmy. Deletions affecting ND5 and ND6 are associated with significantly higher mutant loads, consistent with earlier predictions that disruption of components of a transcriptional feedback mechanism confers a replication advantage^28,29^. This allowed us to estimate *fracAdv* to be approximately 0.14. In contrast, deletions affecting ATP6 are associated with lower heteroplasmy, an unexpected finding that suggests the ATP6 gene product may play a permissive role in processes required for mtDNA propagation, such as transcription or replication. Importantly, such “negative” hotspots, regions where deletions are selectively disfavoured, are largely invisible to classical studies that focus on cells fully dominated by deletion mutants^1-6,8,12-15,17^. In this respect, scRNAseq provides a more comprehensive view of the mtDNA deletion lifecycle, capturing both successful and abortive expansion attempts.

Consistent with both experimental observations and our simulation study^26^, the number of distinct deletion species per cell follows a Poisson distribution with a low mean (approximately 0.1), indicating that most cells harbour no mutants, a small fraction harbour a single mutant, and coexistence of multiple mutant lineages is rare. This strongly constrains mechanistic models and argues against explanations that predict extensive intracellular diversity, such as the vicious cycle hypothesis.

At the same time, the fcwm curves derived from the TMS data deviate from the predictions of the basic Moran model. The proximity of fcwm00 and fcwm05, the threshold-dependent slopes, and the anomalous behaviour of the 3- and 21-month age classes indicate that either data limitations or unmodelled biological processes, or both, affect the inference. Among these, the limited mitochondrial read depth of standard scRNAseq is likely the most severe constraint, as it directly impacts heteroplasmy estimation. Future datasets with targeted mitochondrial enrichment are therefore expected to substantially improve model–data agreement.

Despite these limitations, we proceeded with threshold-specific parameter fits and obtained *selAdv* estimates ranging from 1.57 to 1.7. These values imply mean accumulation times of around 100 days from first occurrence to near-complete dominance. This implies that, once an advantageous deletion arises, clonal expansion proceeds rapidly on the timescale of the murine lifespan. This supports a two-phase view of mtDNA deletion accumulation, indicating a potentially long waiting phase governed by mutation rate, followed by a fast expansion phase driven by selection. Under this view, differences between species in the prevalence of deletion-dominated cells may primarily reflect differences in mutation input rather than fundamentally different expansion kinetics, in agreement with earlier theoretical proposals^28,29^.

Taken together, the present work should be regarded as a proof of concept demonstrating that single-cell cross-sectional omics data can be used to infer key parameters of the mitochondrial lifecycle and to estimate accumulation times of mtDNA deletion mutants. With improved experimental data (like scDNAseq^34-36^) this approach has the potential to provide decisive quantitative tests of competing theories of mitochondrial ageing.

## Methods

### Identification of mtDNA deletions in single-cell RNA sequencing data

To identify and quantify mtDNA deletions in scRNA-seq data, we adapted the software tool MitoSAlt (Mitochondrial Structural Alterations), originally developed by Basu and colleagues for accurate identification of mtDNA deletions and duplications from deep sequencing data^30^. MitoSAlt was designed to address fundamental challenges associated with analysing structural variation in mitochondrial genomes, including their circular topology, high copy number, and frequent heteroplasmy.

A key conceptual feature of MitoSAlt is its explicit treatment of the ambiguity inherent in split-read alignments on circular genomes, where any given split alignment can correspond either to a deletion or to the complementary arc duplication. This ambiguity is resolved by enforcing biologically motivated replication competence constraints, specifically the preservation of essential replication origins, allowing unambiguous and biologically consistent classification of structural events.

The original MitoSAlt workflow consists of three main steps: (i) alignment of sequencing reads to the nuclear and mitochondrial genomes using HISAT2 and LAST, (ii) identification and clustering of split-read breakpoints to define candidate structural rearrangements at single–base pair resolution, and (iii) biological classification of events as deletions or duplications followed by estimation of heteroplasmy and visualization.

Although originally developed for bulk genomic sequencing data, the conceptual framework of MitoSAlt is applicable to transcriptomic reads, which also retain information about underlying mtDNA structure. However, substantial modifications were required to enable analysis at single-cell resolution. We therefore developed a dedicated single-cell–adapted version of the pipeline, termed MitoSAltsc. That version incorporates changes to input handling and data structures to support cell-resolved analysis and was fully reimplemented in Python, replacing the original Perl and R components. This reimplementation substantially improved computational efficiency and scalability, which are essential for processing thousands of individual cells. MitoSAltsc operates directly on raw FASTQ files provided by the Tabula Muris Senis consortium via a public AWS S3 repository (https://registry.opendata.aws/tabula-muris-senis), ensuring reproducibility and avoiding dependence on intermediate alignment products generated by external pipelines. The software is available through our public GitLab repository (https://gitlab.uni-rostock.de/IBIMA/MitoMutantAccTimes2).

The primary output of MitoSAltsc is a cell-resolved catalogue of mtDNA deletion events. For each affected cell, the output includes genomic coordinates of deletion breakpoints and an estimate of heteroplasmy, defined as the ratio of reads supporting the deletion to reads supporting the corresponding wild-type sequence. Cells are identified by unique sample identifiers, allowing direct integration with transcriptomic metadata. In most cases, a single deletion species is detected per cell, but a subset of cells contains multiple distinct deletions with separate heteroplasmy estimates, enabling assessment of intracellular mutant diversity.

## Data and Code Availability

Data and computer code used for this work is available via our GitLab repository at https://gitlab.uni-rostock.de/IBIMA/MitoMutantAccTimes2.

## Acknowledgements

We would very much like to thank Dr. D. Wolz for his help with parameter fitting using fcmaes (https://github.com/dietmarwo/fast-cma-es) and for optimizing the speed of the calculation of mutant distributions based on the Moran process.

## Author Contributions

AK designed the overall concept of the manuscript and developed the mathematical simulations. AK and TK were involved in writing the text and all authors reviewed and approved the manuscript.

## Competing Interests

The authors declare that there are no conflicts of interest.

## Funding Declaration

This research received no specific grant from any funding agency in the public, commercial, or not-for-profit sectors.

## Notes

### Competing Interest Statement

The authors have declared no competing interest.

### Summary of Updates

Figure 5 has been updated. Two additional references have been added.

